# A unifying concept of animal breeding programs

**DOI:** 10.1101/2020.07.08.193201

**Authors:** H. Simianer, A. Ganesan, L. Buettgen, N.T. Ha, T. Pook

## Abstract

Modern animal breeding programs are constantly evolving with advances in breeding theory, biotechnology and genetics. Surprisingly, there seems to be no generally accepted succinct definition of what exactly a breeding program is, neither is there a unified language to describe breeding programs in a comprehensive, unambiguous and reproducible way. In this work, we try to fill this gap by suggesting a general definition of breeding programs that also pertains to cases where genetic progress is not achieved through selection, but e.g. through transgenic technologies, or the aim is not to generate genetic progress, but e.g. to maintain genetic diversity. The key idea of the underlying concept is to represent a breeding program in modular form as a directed graph that is composed of nodes and edges, where nodes represent cohorts of breeding units, usually individuals, and edges represent breeding activities, like ‘selection’ or ‘reproduction’. We claim, that by defining a comprehensive set of nodes and edges it is possible to represent any breeding program of arbitrary complexity by such a graph, which thus comprises a full description of the breeding program. This concept is implemented in a web-based tool (MoBPSweb, available at www.mobps.de) which is described in a companion paper, and has a link to the R-package MoBPS (Modular Breeding Program Simulator) to simulate the described breeding programs. The approach is illustrated by showcasing three different breeding programs of increasing complexity. Finally, potential limitations of the concept are indicated and extensions to other fields, like plant breeding, are discussed.

## Introduction

The book ‘Animal Breeding Plans’ (Lush 1937) is generally considered as the starting point of modern, science based animal breeding, despite the fact that many elements of modern breeding like the fundamentals of genetic inheritance (Mendel 1866), the interplay of variation and selection (Darwin 1859), the concept of quantitative genetics (Galton 1889; Fisher 1918), or the concept of population genetics (Wright 1922) were developed earlier. It was only in Lush’s pioneering book that many of these elements were brought together, giving a grand picture of the complexity of animal breeding and the interplay between the various elements.

Harris *et al.* (1984) later provided a systematic and comprehensive approach for the design of breeding programs. Their system is composed of nine steps, starting with the analysis of the production system and definition of breeding goals and actually covering the entire procedure including the choice of the breeding system and breeds to be used etc.. What we nowadays call a ‘breeding program’ in the narrow sense is basically addressed in their steps five to seven covering (in modern wording) breeding value estimation, selection and mating system. They also address the need to compare alternative variants of a breeding programs with respect to genetic progress and costs.

Definition of breeding objectives addresses the question ‘where to go?’ while breeding strategies describe ‘how to get there’ (Van der Werf in Kinghorn *et al.* (1999)). Such breeding strategies are largely equivalent with what would be called a breeding program in our context and are made up of five major building blocks, namely ‘trait measurement’, ‘estimation of breeding values’, ‘reproductive technologies’, ‘selection & culling’ and ‘mating’. It is further argued that breeding programs are not only targeting to achieve genetic progress in the breeding population, but also dissemination of the genetic progress achieved in a breeding nucleus into the commercial production population must be part of a breeding strategy.

In the same book Kinghorn *et al.* (1999) differentiate between bottom-up and top-down variants of designing breeding programs. In the top-down version, the different steps in the breeding programs are defined and implemented by rules. In the bottom-up version, which he calls ‘tactical implementation of breeding programs’ most beneficial use is made from available animal material and technologies in a somewhat opportunistic sense, in that all available technologies are used and combined in an optimum way. In many real life cases, implemented breeding programs will be a mixture of both approaches, though.

While in the scientific studies discussed above ‘breeding programs’ were - often implicitly - defined in a pragmatic way, there is to our knowledge so far no clear definition, what a breeding program is, nor is there a concept as to how a breeding program can be formally described in a uniform and comprehensive way, i.e. that it reflects all relevant breeding activities in such a way that a third person can fully comprehend and reconstruct the respective breeding program from description alone.

However, such a formal definition would be very useful for many reasons: It requires a clear specification of the relevant parts and processes of a breeding program and an explicit delimitation of its scope, i.e. at which point does a breeding program start, at which point does it end, and which elements and activities are seen as parts of the breeding program and which ones are not. It also provides the basis for the description of a breeding programs in a unified structure and terminology, allowing e.g. to formally compare different versions of a breeding program. Finally, it can be used as a basis to assess outcomes of a breeding program in a quantitative sense. These can range from rather simple quantities like the duration of a breeding cycle or the costs of a breeding program to rather complex, but highly relevant parameters such as genetic progress or the development of inbreeding. The latter would require suitable deterministic or stochastic analysis tools that can read in the respective breeding program description and calculate the relevant quantities. This obviously would require a highly formalised ‘breeding program description language’.

Interestingly, there is a legal definition in the “REGULATION (EU) 2016/1012 OF THE EUROPEAN PARLIAMENT AND OF THE COUNCIL of 8 June 2016 on zootechnical and genealogical conditions for the breeding, trade in and entry into the Union of purebred breeding animals, hybrid breeding pigs and the germinal products thereof”, stating: “Breeding program means a set of systematic actions, including recording, selection, breeding and exchange of breeding animals and their germinal products, designed and implemented to preserve or enhance desired phenotypic and/or genotypic characteristics in the target breeding population.” Actually, this is a very thoughtful definition which largely is in agreement with our understanding of what a breeding program is.

A key requirement for a formal assessment of breeding programs is that one is able to predict genetic progress achieved through this program. The so-called “breeders’ equation” (Lush 1937) provides a simple prediction for the short term genetic response when selecting in a homogeneous population. However, most breeding programs are not conducted in entirely homogeneous populations but rather are stratified into clusters of breeding animals that differ in relevant characteristics, like sex, information on own or relatives’ performance, reproduction rate, or cluster size and, through this, selection intensity. This conceptually was first reflected in the four-path model of Rendel and Robertson (1950) which allows for different selection parameters (like accuracy of estimated breeding values, selection intensity, and generation interval) on the pathways leading from the four grand-parents to the grand-children. But it was soon realized that most breeding programs are even more complex, especially when generations are not discrete, but overlapping, as they are in many real-life situations. With the introduction of the so-called ‘gene-flow’ approach by Hill (1974); Elsen and Mocquot (1976) it became possible to assess expected genetic progress from breeding schemes of arbitrary complexity. In this concept, groups of animals of the same sex, age, and selection background (often called ‘cohorts’) are defined as units of a breeding program and transition matrices are used to model the gene-flow between such groups through aging and reproduction in discrete time steps. A system based on matrix algebra then allows to derive the expected genetic progress. Later, deterministic breeding program simulations like ZPLAN (Nitter and Graser 1994) and ZPLAN+ (Täubert *et al.* 2010) made this concept available for the assessment of complex breeding schemes.

While the fundamental concept of the gene-flow approach, namely that breeding programs can be basically represented by cohorts and the gene flow between them, was a major break-through, it has some limitations in reflecting the true complexity of modern breeding programs and providing all relevant information. Firstly, the gene-flow concept intrinsically assumes a quantitative genetic background of traits under selection and does not naturally provide an extension to account for traits that are exclusively or partially inherited via major genes. Also, genetic progress conceptually may not be generated through selection only, but also through other means like gene editing (Jenko *et al.* 2015). It is also conceptually difficult to represent practically relevant breeding practices other than selection, e.g. nonrandom mating (Liu *et al.* 2017), in such a concept. Secondly, while genetic progress is of major importance in the assessment of breeding programs, other parameters, like changes in allele or genotype frequencies at certain loci or the development of relationship and inbreeding often are relevant as well. Although deterministic solutions for some of these problems exist (Bijma *et al.* 2001), these are rather complex and have limitations when applied in a recursive manner.

With rising computational power, stochastic simulation have become viable alternative to deterministic simulation. Simulators such as AlphaSim (Faux *et al.* 2016) and MoBPS (Pook *et al.* 2020b) provide simulation frameworks that are simulating individual meioses, generate genotypes and phenotypes for single animals or allow to specifically select which individuals to mate. Based on this, more complex breeding programs can be simulated and compared against each other. The included stochasticity is both a blessing and a curse, as the variability of outcomes can be analyzed, but a high number of simulations is required as, in contrast to deterministic simulations, no expected values are obtained. Besides breeding, stochastic simulation has also shown to be highly useful for a variety of applications in population genetics with simulators such as forqs (Kessner and Novembre 2014), XSim (Cheng *et al.* 2015), SLiM (Messer 2013), QMSim (Sargolzaei and Schenkel 2009).

The main objective of a breeding program is to generate genetic progress, which will lead to a more economical production as well as to improvements in other areas, like animal welfare or environmental impact. In general, the economic benefit of the achieved genetic improvement is difficult to quantify, since the potential to generate monetary returns depends to a large extent on factors outside the breeding program itself, like consumer preference or competition of different breeding programs in the market (Dekkers and Shook 1990). Also, it needs to be defined for which level the economic benefit should be derived. Economic benefits through genetic gains can be calculated for a national economy, for a sector (like agriculture or food production), or for different institutional actors in multi-actor breeding programs. In a dairy cattle breeding program these can e.g. be single farms, breeding organizations or AI companies. Note that the economic interests of these actors are not necessarily congruent: while a farm wants to purchase cheap semen, the AI company aims at selling semen for a high price.

Another important aspect is that genetic gains are cumulative and ‘eternal’, i.e. once a genetic population mean is raised to a higher level through breeding, this level will sustain without any further investment (Weller 1994). Since relevant genetic progress occurs only after some generation intervals, the time horizon for the economic evaluation of genetic gains typically is long and costs and returns need to be discounted for a fair appraisal. Hill (1971) suggested an approach to calculate cumulative discounted returns of a breeding investment over an extend investment period which takes these factors into account. This concept for the economic evaluation of breeding programs was implemented in the ZPLAN software (Nitter and Graser 1994) and used e.g. for the economic evaluation of genomic dairy cattle breeding programs (König *et al.* 2009).

Conservation breeding programs, aiming at maintenance of genetic diversity, also suffer from imbalances with respect to stakeholders and time scale: the concrete costs have to be borne today by the breeding program, while the benefit will be hard to quantify on a monetary basis, may occur only with some probability at an unspecified time point in the future, and will be on a national or even global scale, as argued by Smith (FAO 1984) and Simianer (2005).

What is quantifiable, though, are the actual costs and returns occurring in the breeding program itself. Accountable costs are all costs that are directly caused by the breeding program, such as expenses for performance testing, genotyping, data recording, or breeding value estimation, where costs can be either fixed or variable, i.e. depending on the number of individuals. Accountable returns can arise if e.g. animals or products from a performance test are sold. For all costs and returns it is essential that they appear at an identifiable time point in a breeding program. Since a reasonable evaluation of breeding programs makes only sense in a medium or long term, usually stretching over decades, it is essential to consolidate discounted expenses and returns on the same time point. With this it will be possible to compare the consolidated and discounted costs of a breeding program with the expected genetic gain.

In the following we first give a general definition of what we consider as a breeding program and refer to some implications, making reference to practical breeding. Later we provide a formal framework to describe a breeding program in a modular way and discuss a possible set of modules of a breeding program without claiming this set to be comprehensive. This modular concept was implemented in an interactive software tool, the graphical user interface of the Modular Breeding Program Simulator (Pook *et al.* 2020a) which will be introduced and illustrated with a few examples of increasing complexity. Finally, the usefulness of the concept and potential extensions to other fields of application will be discussed.

## Methods

### What is a breeding program?

We define a breeding program as a structured, man-driven process in time that starts with a group of individuals *X* at time *t*_1_ and leads to a group of individuals *Y* at time *t*_2_ *> t*_1_. The objective of a breeding program is to transform the genetic characteristics of group *X* to group *Y* in a desired direction, and a breeding program is characterised by the fact, that the implemented actions aim at achieving this transformation. To stay as general as possible we deliberately avoid terms like “selection” or “genetic progress”, since the goal of a breeding program may also be in other fields than genetic gain, e.g. it can be maintenance of genetic diversity, and goals can also be reached by means other than selection, such as genetic engineering.

All groups of individuals and all activities that contribute to this transformation process are considered as elements of the breeding program. This includes direct actions and groups the breeding activities act on, but also groups of individuals that provide information which is used in the transformation process. This means, that in a nucleus breeding program the individuals generated in multiplier herds are considered as part of the breeding program, if their performance data are used for selection decisions in the breeding nucleus, but are not part of the breeding program if this information is not used for a breeding value estimation or selection. A simple possibility to find out if a group of individuals must be considered as part of the breeding program or not is to ask, whether the absence of the group would affect any of the breeding activities in a direct or indirect way.

This definition also means, that important activities in the breeding process, like defining the breeding objective, choosing the populations to work with, estimation of economic parameters etc. are not part of the breeding program as we define it. Also choosing or optimizing the design of a breeding program is not considered as part of the breeding program itself, but would be a result of comparing different breeding programs, e.g. based on comparative quantitative assessments of outcomes resulting from alternative breeding program designs.

Breeding programs are stochastic processes by nature in that certain parts have an implicit randomness. This is the case for Mendelian sampling when forming gametes during meiosis, but also for phenomena like mortality during aging or success rate of certain reproduction techniques. Here we assume that a certain proportion of individuals dies at random up to a certain age, or that a reproduction technology is only successful in a subset of individuals, where in both cases the decision on which individual is affected is completely random. If it is not completely random but a genetic component is involved, there are possibilities to model this as part of a breeding program, though.

We further assume breeding programs to take place in a constant environment or set of environments, that is we assume all environmental factors to be invariable over time. By this, all observed genetic or phenotypic changes can be attributed directly to the breeding program (taking into account the intrinsic stochasticity of the involved processes). This concept deviates from the real life situation, where it is often seen that genetic improvements go along with improvements in the production conditions and thus the observed phenotypic trend can be attributed in parts to breeding and to environment, respectively (for an impressive example see Havenstein *et al.* (2003)). The constant environment assumption also encompasses the biotechnologies used in breeding, like artificial insemination or embryo transfer, which are assumed to have constant success rate over the considered time span.

### Formal description of a breeding program

For a formal description of a breeding program, three characteristics are essential:

- Comprehensiveness: The description should contain all relevant elements and steps in the breeding program
- Unambiguousness: The description should be fully understandable to a breeder by using a well defined terminology
- Reproducibility: From the description alone it should be possible for a breeder to reconstruct the described breeding program in all relevant aspects

To fully describe a breeding program, two parts are needed: the breeding environment and the structure of the breeding program.

The breeding environment includes the definition of the founder population(s) to work with, traits that are relevant in the context of performance testing, breeding value estimation, and are included in the breeding goal (note, that these are not necessarily identical sets of traits). For the joint set of traits we need to define all required parameters (phenotypic mean and variance in the founder population, heritabilities, repeatability, genetic and phenotypic correlations). In general, traits can be considered either to have a quantitative genetic background, meaning that genetic variances and covariances are defined explicitly, or to have a discrete genetic background. In this case, traits are assumed to be affected by a set of hypothetical genes and variances and covariances result from the assumed genotype frequencies, direct and pleiotropic genotype effects at the assumed loci and linkage disequilibria between them. In both ways complex genetic structures like dominance and epistasis as well as a transformation from a quantitative background to a discrete phenotype via a logistic model can be included. In case a genomic breeding program is to be modelled, it is essential to define the type of genomic information (e.g. the assumed SNP array or marker density used) that is available on the genotyped individuals.

If the objective of the breeding program is to make genetic progress we also need to define the breeding goal, which typically can be implicitly formulated in an economic form by providing marginal economic values per natural unit or genetic standard deviation of each trait included in the breeding goal. If the breeding goal is more complex, e.g. by putting a restriction on a certain trait or other variables, like maintenance of a certain diversity level, this also must be stated.

In many breeding programs, there are typical phenotyping and selection scenarios, i.e. for a certain group of individuals a specific set of traits is recorded, which may depend on sex, age, environment (e.g. field vs. station testing) and other factors. Also, typical combinations of data (own or relatives’ phenotypes, genotypes etc.) are available at certain stages of a breeding program. This, together with the type of breeding value estimation (e.g. selection index, pedigree-based BLUP, various types of genomic predictions) applied at a given stage need to be defined as well.

Once the breeding environment has been defined, the structure of the breeding program can be represented as a directed graph, being composed of nodes, which are made up of elementary objects, and edges, which link nodes in a directional way. The three key items are defined as follows:

An elementary object is the smallest unit to be addressed in a breeding program. Elementary objects exist in different types, which e.g. can be ‘individual’ or ‘gamete’. A given type has a certain set of attributes (e.g. an elementary object of class ‘individual’ can have the attributes ‘sex’, ‘age’, ‘phenotype’, etc.). A node is a group of elementary objects which all belong to the same type. Nodes can have two given attributes: the node size which is a positive integer, and the duration of the node, which reflects the time allocated to the ‘lifespan’ of this node on an appropriate time scale. A node can be either a founder node, which has no predecessors, or a derived node which results from one or several other nodes through edges. We do not allow for ‘partial’ founder nodes, which would e.g. be the case if an offspring node is generated from a known paternal node but the maternal origin is unspecified.

An edge is a directed link from one node of origin to one target node. Edges exist in different classes which e.g. can be ‘aging’, ‘selection’ or ‘reproduction’. A given edge class has a certain set of attributes (e.g. the edge ‘selection’ can have the attribute ‘selected proportion’, and ‘selection type’, which can be ‘random’, ‘phenotypic’ etc.). An edge of a specific class defines the transformation rule of the set of elementary objects in the node of origin into the set of elementary objects in the target node. The transformation rule is applied to all elementary objects of the node of origin in a uniform, possibly probabilistic way, so it does not systematically discriminate between the elementary objects in that node. Direction is forward (never backward) in time, therefore no loops are possible. Edges can also be given a certain time needed to be executed, e.g. a ‘Reproduction’ edge in dairy cattle typically should have a nine month time span.

Next, we present various types of elementary objects, nodes and edges that we consider useful in describing relevant breeding programs. We do not claim that the presented lists of items are entirely comprehensive, but they should be sufficient to model most of the contemporary animal breeding programs. If necessary (e.g. due to future technological progress) the list can be easily extended. Possible classes of ‘Elementary objects’ are:

- Individual
- Gamete
- Gene

With ‘Individual’ certainly being the most important class of elementary objects. Each individual, as each elementary object in general, must have a unique identifier so it can be traced through the breeding program (note that the same elementary object may occur in several nodes). Key attributes are sire, dam, phenotype, true genomic value, and estimated breeding value. In many cases a genotype at a predefined set of loci will be useful. This can encompass everything between a genotype at a single locus and a whole genome sequence, including practically highly relevant cases as e.g. genotypes for standard SNP arrays. Individuals further can have ‘age’ as an attribute in case ageing is of relevance and animals have different age-dependent properties (such as repeated performances like first, second and third lactation). Individuals thus can have different performance traits which can be dependent on sex, age, and possibly other factors. For individuals in cohorts undergoing selection it is essential to provide a variable allowing to rank all members of the cohort. This can be as simple as a phenotype, but in many cases will be an estimated breeding value which may incorporate information not only from the respective cohort, but from additional past and contemporary cohorts as well. Since they have a genetic makeup which is identical to a living animal, embryos in most cases can be modelled as a special type of individuals.

‘Gamete’ is a second potentially relevant class of elementary objects and can be either male (sperm) or female (oocyte). Gametes carry a haploid genome and a gametic value. It can make sense to define them explicitly in a breeding program if they are targets of biotechnological manipulations, like e.g. sperm sexing or transgenic technologies.

In some applications it will make sense to consider also ‘Gene’ as an elementary object, e.g. if genes are targets of manipulation (e.g. gene editing) or they are to be transferred into an existent genome (e.g. through marker-assisted introgression or technological gene transfer).

As said before, a node is defined as a group of elementary objects, which all belong to the same class. The minimum group size is one, allowing to include a specific individual with certain characteristics (e.g. the carrier of a specific mutation) in the design of the breeding program. Note that elementary objects in the same node, while being of the same class, may have different attributes, which means that an attribute can be present in a subgroup of the elementary objects in a node and lacking (i.e. NA) in another subgroup. However, attributes will usually be similar as the same breeding actions are applied upon all elementary objects of a node. In case all elementary objects in a node have the same value for a given attribute, we will refer to this as the attribute of the node.

Some of the more important ‘Edge’ classes, with no claim of

- completeness, are the following:
- Aging
- Selection
- Reproduction
- Split
- Combine
- Cloning
- Repeat

The edge class ‘Aging’ links two cohorts where the individuals in the target cohort are identical to or a subset of the individuals in the cohort of origin, while some characteristics, especially ‘age’, but in most cases also ‘phenotype’ change. Aging transfers the individuals to a later age, which preferably should be defined on an appropriate time scale (weeks, months, years), but also functional time scales (first, second, third lactation) can be used. If aging goes along with a decrease in the node size, the individuals to appear in the target node are a random subset of the individuals in the node of origin, reflecting natural mortality or other factors, like animal sales. In case dropping out appears not at random, but is associated with fitness, a corresponding fitness trait should be modelled explicitly and one should rather use an edge of class ‘selection’ than ‘aging’.

‘Selection’ is similar to aging, but it encompasses a systematic choice of the individuals to be moved to the target node. Typically, individuals in the parent node are ordered in descending order (from best to worst) for a single variable (which can be a phenotype or an estimated breeding value, but also a random number) and only the top proportion is moved to the target node. While an edge is exclusively using the information in the node of origin to derive the properties of the elements in the target node, attributes of the elementary objects in a node may include ‚collapsed’ information that is based on other past or contemporary nodes, as e.g. estimated breeding values which are calculated from all performance and pedigree data available at the given time point.

The edge class ‘Reproduction’ is needed to generate individuals of the next generation. This edge class typically occurs in pairs, linking one or more paternal and a maternal nodes, to a single offspring node. The underlying transformation comprises the formation of a gamete on either parental side, choosing at random one individual of the respective parental node and creating a gamete by random Mendelian sampling. The two parental gametes are then combined, forming the genotype of the offspring. It is further necessary to define the frequency of the use of the individuals in the parental node, which can e.g. be random or uniform or as a function of the genomic value, and possibly whether mating appears at random or follows a pattern, e.g. assortative or disassortative mating according to a defined ranking criterion.

Further useful edge classes are ‘Split’ and ‘Combine’. ‘Split’ creates two or more nodes out of one by assigning the members of the node of origin to respective target nodes either at random or according to a defined criterion. This e.g. is useful to generate different mating classes like bull dams and cow dams. ‘Combine’ merges two or more nodes into one, which e.g. can be useful in a case of overlapping generations when the best animals are to be selected from several generations in a competitive setting. Then, the available animals can be combined in one node and the best ones from this node can be selected based on estimated breeding values. Additional edges can be defined when respective biotechnologies are used, such as ‘Cloning’, which allows to genetically duplicate existing individuals, or ‘Editing’ reflecting the possibility to modify the genotype of an individual in a targeted way. With more technologies coming up more specific edge classes may become necessary. Objects of such an edit can be nodes of class ‘Individual’, ‘Gamete’, or ‘Gene’.

A useful edge class to describe breeding programs that are composed of several breeding cycles is ‘Repeat’. It can be used to copy resulting nodes from one breeding cycle into the nodes of origin of the next cycle, assuming that exactly the same breeding activities are to be repeated in each cycle. The ‘repeat’ edge has the attribute ‘number of repeats’ which allows to determine the desired number of cycles.

With this combination of the breeding environment together with the limited set of elementary objects, nodes and edges with their respective attributes, we claim that it is possible to describe any animal breeding program of arbitrary complexity in a comprehensive, unambiguous and reproducible way. The only additional information required for a full documentation is to determine which data are used to calculate estimated breeding values at which stage. It should be evident that only performances that have been completed prior to the point of time where the breeding value estimation is conducted can be used, but in many cases only a subset of conceptually available data is actually used on different selection paths, which needs to be specified.

This general concept was implemented in a web-based graphical user interface MoBPSweb (Pook *et al.* 2020a) which is accessible under www.mobps.de. This interface offers a wide range of options to define breeding programs along the concept outlined above and subsequently simulate the breeding program with the MoBPS R-package (Pook *et al.* 2020b) as the backend simulator. Nodes and edges are defined in a very general way, and together can form breeding programs of almost arbitrary complexity. Templates are provided for standard cases. Once a breeding program is described, it can be exported in form of a flat file in JavaScript Object Notation (JSON) format, which contains all relevant information and thus can be seen as a full description of the breeding program. This file can be archived and later imported again, displaying the full breeding structure without any loss of information.

### Time scale, resource allocation and economics

Each edge and node can be assigned a duration in time units on an appropriately chosen scale (weeks, months, or years) and should reflect the time needed for the respective breeding activity in an edge or the time a certain node is existing. To provide a realistic outcome, realistic durations should be assigned here. These then can be used to give a time stamp to each node, reflecting the earliest possible time point when such a node comes into existence, which is the maximum sum of durations of edges and nodes along all possible paths from the founder nodes, which are assumed to be contemporary and have the initial time stamp ‘zero’.

From the node-and-edge representation of the breeding program, the required resources can be extracted and can be displayed along the time scale. This encompasses e.g. the occupied testing capacity, the number of animals to be genotyped, or the time points when breeding value estimation needs to be conducted together with the expected number of individuals and performance records to be processed. It is also possible to derive the development of costs …

### Data Availability

All breeding programs described in this manuscript can be found as templates in the MoBPSweb (Pook *et al.* 2020a) at www.mobps.de. Corresponding JSON-files of the breeding programs can be found in Supplementary Files S1, S2, S3. Supplementary files are available at FigShare.

## Results

### Illustration of the concept with different case studies

In the following, we will first give a very basic example for a breeding program in a very detailed form, and later we present more complex breeding programs with less detail to illustrate the main idea of the concept, building upon clear definitions of elementary objects, nodes and edges, which can be flexibly combined in order to represent breeding programs of arbitrary complexity. All breeding schemes were entered, evaluated and simulated via the graphical interface (available at mobps.de) with the R-package MoBPS (Pook *et al.* 2020b) as the backend simulator.

### A basic breeding program

In the following, we define the breeding environment as well as a minimum set of elementary objects, nodes and edges that is sufficient to define a basic breeding program.

As the breeding environment we define to start with a noninbred and unrelated random set of simulated individuals as a founder population. We consider a single quantitative trait, which can be measured in both sexes. We aim at increasing the phenotypic level for the considered trait, and selection is on an individual’s own performance. For a backend simulation as done in MoBPS (Pook *et al.* 2020b) one would typically store underlying genomic data and assume the presence of underlying QTL leading to *𝒩*(*µ, σ*^2^) distributed traits with a given heritability.

For this we just need a single class of elementary objects, namely the type ‘individual’, with several different attributes (Table 1). The nodes are then generated by combining all / the selected elementary objects (individuals) in a joined cohort. Finally, we need three edge classes, namely ‘Selection’,’Reproduction’ and ‘Repeat’ (Table 2).

**Table 1.** Elementary objects for the basic breeding program.

| class | attributes | possible values |
| --- | --- | --- |
| individual | sex | 'male'; 'female' |
|  | phenotype | real number; NA |
|  | genotype | marker sequence; NA |
|  | father | an elementary object of class 'individual'; NA |
|  | mother | an elementary object of class 'individual'; NA |

**Table 2.** Classes of edges used in the basic breeding program.

| class | attributes | possible values |
| --- | --- | --- |
| selection | node of origin |  |
|  | target node |  |
| | selected proportion | real number $0 < \pi \leq 1$ |
|  | selection type | 'random', 'phenotypic' 'true genomic value', 'estimated breeding value' |
| reproduction | node of origin |  |
|  | target node |  |
| repeat | node of origin |  |
|  | target node |  |
|  | number of cycles | positive integer |

For the edge class ‘reproduction’ we need to define the additional requirement that it is only admissible in pairs, meaning a second edge of class ‘reproduction’ must link the chosen target node with a node of origin of the other sex.

An elementary breeding program constructed with this basic set of nodes and edges is displayed in Figure 1. Here, selection is both in males and females, however with different selection intensities, and the shown breeding cycle is repeated 10 times. Details on the attributes of all nodes and edges of the breeding program can be found at Supplementary File S1.

**Figure 1.**
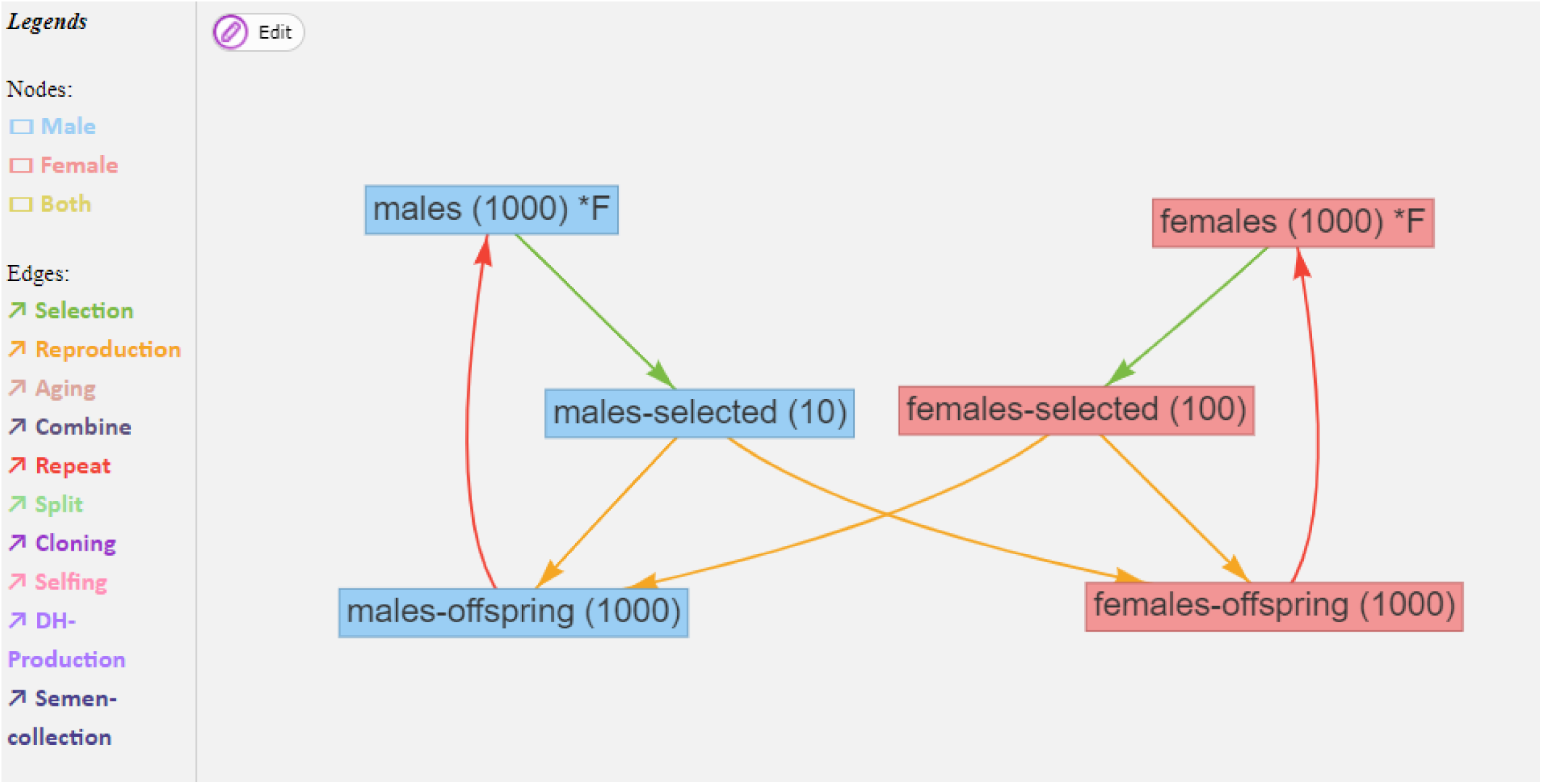
An elementary breeding program. Nodes in blue (red) depict male (female) cohorts, with number of individuals given in brackets. Green, orange, and red edges represent selection, reproduction, and repeats, respectively. Initial founders of the breeding scheme are indicated by “*F”.

### Dairy cattle farm

As a second example, we consider selection in a dairy cattle on farm level (Figure 2). Animals are split based on their current age resulting in five different age groups at each time period (calf, heifer, cow-L1, cow-L2, cow-L3, an extension to higher lactations would be straightforward). With the exception of calves which are too young to reproduce, all individuals are potentially used for reproduction of the next generation of calves (calf+). The other cohorts of the next time period are generated by selecting from the respective one level younger cohort of the current time period. We here consider multiple traits (milk, fertility, health traits) of which some are only observed in female individuals and/or in certain age cohorts. Furthermore, phenotypic information potentially accumulates over time (e.g. milk yield from repeated lactations) and information selection is based on becomes more accurate. The paternal breeding material is introduced in the form of semen from a breeding company.

**Figure 2.**
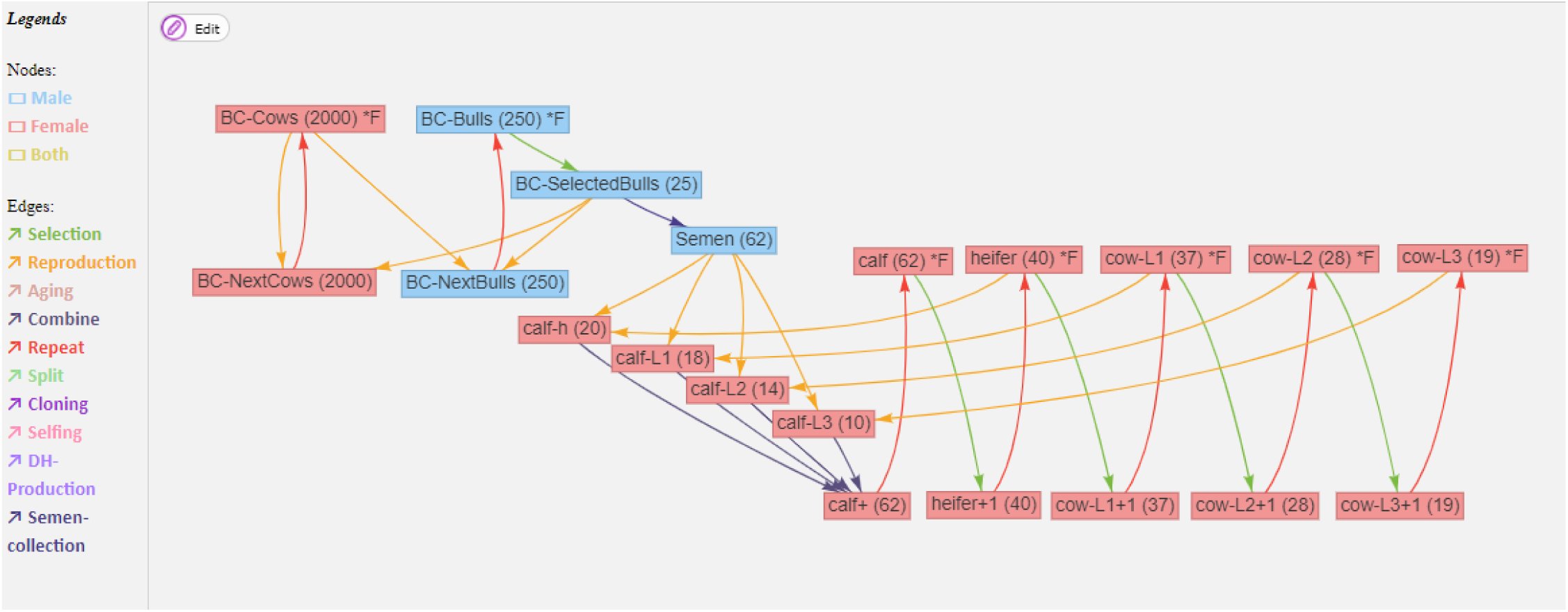
A dairy cattle selection program within a herd, with animals separated in age groups. Details on the attributes of all nodes and edges can be found in Supplementary File S2.

With rising complexity of the breeding program the required set of attributes for individuals, nodes and edges is increasing. E.g. for each individual it is now necessary to track if the respective individual is genotyped, and which phenotypes are available at the time of selection for the animal itself and its relatives. As for the nodes, the time point of generation needs to be tracked to infer the current age of all individuals in the respective node. In regard to edges the two classes “Combine” and “Semen-collection” need to be added. Furthermore, additional options like the method used for breeding value estimation (“genomic BLUP”, “pedigree BLUP”, “single step BLUP”, “BayesA” etc.) and the cohorts that are used in the breeding value estimation need to be provided. Extended discussion on this particular breeding program and a variety of advanced scenarios (e.g. change of weights in the selection index, reducing the selection intensity on the paternal side) can be found in Pook *et al.* (2020a).

### Commercial layer breeding program

As a second example, we consider a commercial layer breeding program with a four-line cross as described in Sitzenstock *et al.* (2013). The scheme for one breeding cycle in line A, one of the four lines, is given in Figure 3. To keep an overview, we used a systematic nomenclature of nodes representing line-sex-housing-age-breedingstatus with the different options described in Table 3. In general, selection of cocks and hens is modeled on the right side of Figure 3. Information is generated from the female selection candidates in single and group cages, but also from massive numbers of full- and halfsibs tested in single- or group cages (center of Figure 3), as well as from paternal halfsibs which are crossbreds with line

**Table 3.** Nomenclature for the commercial layer breeding program given in Figure 3

| Type | Options |
| --- | --- |
| Line | A, B, C, D |
| Sex | M (male), F (female) |
| Housing | PE (practical environment), SC (single cage), GC (group cage) |
| Age | 0, 51, 91 (in weeks) |
| Breeding status | Test (phenotypic testing), Cand (selection candidate), Sel (selected individuals) |

**Figure 3.**
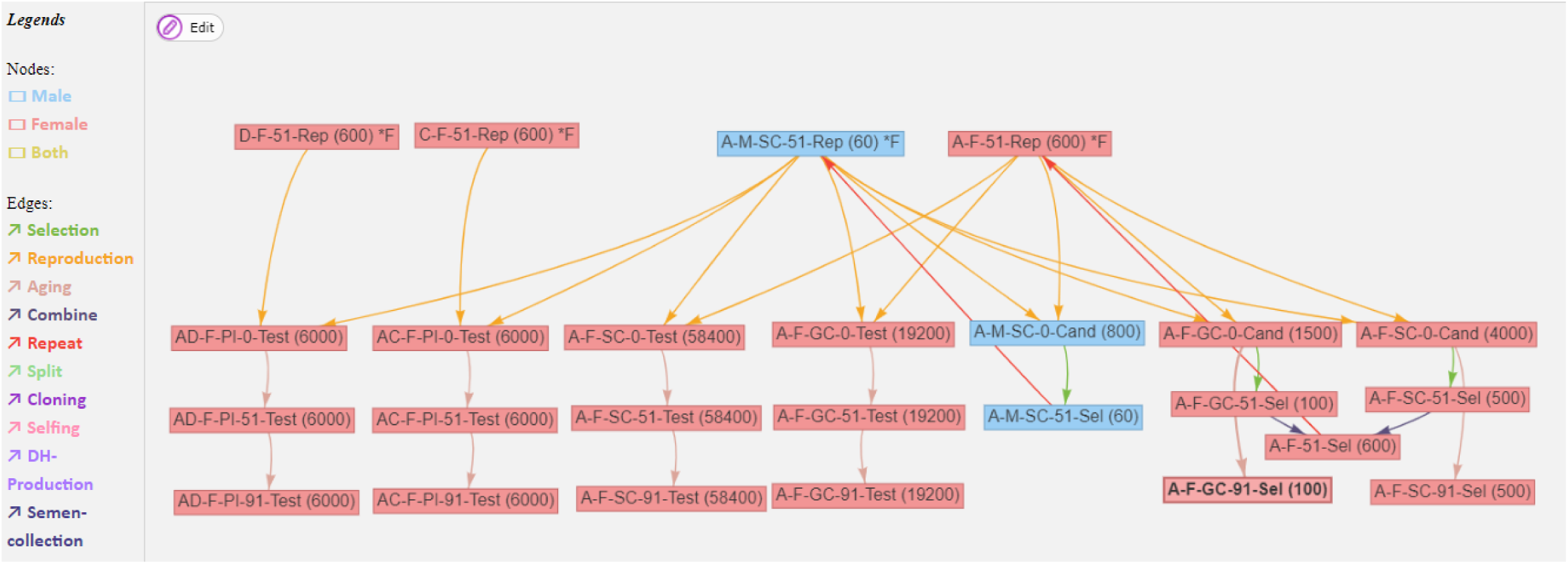
Breeding scheme for line A of a four-line cross in a commercial layer breeding program as described in Sitzenstock *et al.* (2013). Details on the attributes of all nodes and edges can be found in Supplementary File S3.

C and D, respectively (left side of Figure 3). We further consider three ages: 0 weeks when chicks enter the breeding program, 51 weeks, when selection of hens and cocks is performed, and 91 weeks, when for all animals the entire lifetime performance is available, which eventually will be used as information in the breeding value estimation in subsequent breeding cycles. Traits considered in this breeding scheme are egg weight, feed consumption, egg-shell strength, mortality and laying performance in different laying periods, and for each node it is specified which set of traits is recorded for the respective animals in that node. Variable costs can be attributed to rearing (housing), performance testing (phenotyping) and genotyping. The interested reader is referred to Sitzenstock *et al.* (2013) for details on the cost structure of the breeding program. Details on the attributes of all nodes and edges of the breeding program can be found at Supplementary File S3.

## Discussion

The suggested concept aims at describing breeding programs in a standardized, unambiguous and reproducible way. This is an essential prerequisite for a number of useful applications:

It provides the possibility for a clear description and fully standardized documentation of a breeding program. Today, breeding programs are usually described in an informal verbal way, using terminology that often is not clearly defined (see e.g. Mueller *et al.* (2015)). Such descriptions often do not fulfill the criteria of comprehensiveness, unambiguousness and reproducibility as defined earlier, since, e.g. not all elements of the breeding program are sufficiently described, or terminology may be used in an ambiguous manner. This may cause the communication about a breeding program design in a breeding company to be error prone, possibly leading to deviations between the intended design and the actual implementation. It should also be seen, that outcomes of breeding programs generally materialize with a considerable time lag, and it may happen that the breeding program that has led to the actual genetic progress is not well enough documented, making it hard to trace back current results to possible causes a posteriori.

The standardised description of a breeding program presented here and implemented in MoBPSweb (Pook *et al.* 2020a) opens various possibilities. Part 2 of Annex 1 of the “REGULATION (EU) 2016/1012 OF THE EUROPEAN PARLIAMENT AND OF THE COUNCIL of 8 June 2016” describes the “Requirements for the approval of breeding programs carried out by breed societies and breeding operations”, part of which is a detailed description of the breeding program to be implemented. So far each breeding association has to find it’s own way of describing the breeding program, and on the other side the authorities are faced with reading and reviewing a variety of possibilities to describe breeding programs. The use of a formalized description of a breeding program would make this process easier and more efficient for both sides.

As shown in the examples, a standardized documentation of a breeding program can be considered as the core of a management tool for breeding programs, which allows to plan the costs, time and resources required to implement the program. Essential information that can be extracted from such a formalized description of a breeding program is a detailed planning for the use of resources, which e.g. comprises a time grid for the number of testing units for phenotyping, the batch sizes for genotyping in genomic breeding programs, a plan at which time which type of breeding value estimation needs to be performed for which population size, etc..

Finally, a formal breeding program description can be used as input for a software to simulate breeding programs. In fact, the JSON file to be exported from MoBPSweb (Pook *et al.* 2020a) can be used as an input parameter file in the R-package MoBPS (Pook *et al.* 2020b), which allows stochastic simulation of the described breeding program (json.simulation()). Such a stochastic simulation software not only provides an estimate of the genetic trend over time, but also relevant parameters like the development of inbreeding and relationship, the development of allele frequency at certain loci of interest, etc.. A further layer, which is yet to be developed, may even allow for the optimization of such breeding programs, e.g. based on finding the optimum allocation of limited resources in a predefined general scheme with variable elements (such as the number of animals in performance testing or the number of genotyped selection candidates).

The described concept has only few limitations, since the modular architecture allows to describe breeding programs at any level of complexity. Eventually newly arising breeding technologies can be implemented easily by defining and adding novel types of edges to the system. In some cases, the concept may come to its limits, though. Such a case might be if the number and structure of nodes is not fixed, but is determined as an outcome of the breeding program itself. As an example, consider stochastic effects that affect the number of individuals in each cohort. The simplest case of this being the sex determination of an offspring or, more generally, any inheritance pattern with Medelian sampling, as well as processes showing variability due to technical or organisational reasons. It is well known, that success rates of reproduction technologies in a breeding context, like artificial insemination or embryo transfer, show considerable variability (König *et al.* 2007; Samper and Morris 1998). Such biological variability may immediately cause a delay in reproduction by one or more reproductive cycles, which in turn will cause fluctuation in generation intervals (Wallinga and Lipsitch 2007). If not strictly controlled for and managed, such a variability will accumulate over repeated breeding cycles and lead to overlapping time patterns which can only imperfectly be represented by a fixed breeding scheme.

This in turn would lead to a change of the selection intensity and/or the number of individuals in subsequent steps or in the most extreme case even lead to entirely different breeding actions being performed in later steps.

Despite such limitations, it should be seen that modeling always has a simplifying component, or, to use a famous quotation, “all models are wrong, but some are useful” (Box 1976). To provide useful insights it is essential that a model extracts the most relevant patterns, while one should always have in mind that implementations are affected by stochasticity and thus may deviate from the simplified structures captured by the model.

While the concept was developed and described in the context of livestock breeding programs, extensions to other fields where design of programs for genetic improvement or genetic management is relevant. This obviously is the case in crop breeding programs (Smith *et al.* 2006), where the basic structure and the objectives are similar to livestock breeding, but differences arise in many details. Major differences lie in the fact, that animal breeding generally is based on individual phenotypes, while in plant breeding averages over many individuals (often in replicated plots) are used, a variety of natural or technologically induced reproductive patterns are available (such as self-pollination or doubled haploid techniques), and most plant breeding program are targeted to achieve a consolidated and uniform variety, while most animal breeding programs aim at a continuous and permanent improvement in a heterogeneous breeding population. But still an extension of the concept to be applicable in a plant breeding context is rather straightforward and will require, among other things, the definition of plant-specific reproduction edges and multiple-individual nodes in performance testing.

Another potential area of application of the concept would be in breeding and management of model organisms in a research context. Among the widely known model organisms are the mammalian model organisms mouse and rat, zebra fish as a vertebrate, Drosophila melanogaster as an invertebrate, plant models such as Arabidopsis thaliana and poplar, or yeast as a fungus, however the actual list is much longer and continuously expanding. Model species breeding (Chesler *et al.* 2008) usually does not aim at genetic improvement, but at the development of genetically homogeneous stocks with defined genetic characteristics (e.g. carrying a certain genotype or being a knock-out for a defined genotype). For this, methods of classical breeding are often combined with transgenic technologies. Again, only minor modifications and extensions would be required to adapt the method to this field of application.

A further potential field of application may be in modelling of diversity management of natural populations. An often-studies case in this context is the genetic dynamics of a subdivided population in discretized habitats (Heino and Hanski 2001) which can be represented in a network graph framework with nodes and edges (Greenbaum and Fefferman 2017). Our approach could be easily extended to model such patterns by defining discrete local subpopulations with migration edges between those nodes which are reflecting physical conditions, like geographic distance or topological conditions along the connecting corridor.

## Acknowledgments

This study was conducted in the context of the European Union’s Horizon 2020 Research and Innovation Program under grant agreement n°677353 IMAGE.

